# Microstructural properties of the vertical occipital fasciculus explain the variability in human stereoacuity

**DOI:** 10.1101/288753

**Authors:** Hiroki Oishi, Hiromasa Takemura, Shuntaro C. Aoki, Ichiro Fujita, Kaoru Amano

**Affiliations:** Graduate School of Frontier Biosciences, Osaka University, Suita 565-0871, Japan; Center for Information and Neural Network (CiNet), National Institute of Information and Communications Technology, and Osaka University, Suita 565-0871, Japan

**Author notes:** These authors contributed equally (Hiroki Oishi, Hiromasa Takemura). **Author contributions** H.O., H.T., I.F., and K.A. designed research; H.O., and H.T. performed research; H.T., and S.C.A. contributed new reagents/analytic tools; H.O., and H.T. analyzed data; H.O., H.T., I.F., and K.A. wrote the paper.

**Keywords:** diffusion MRI, vertical occipital fasciculus, stereoacuity, white matter

## Abstract

Stereopsis is a fundamental visual function that has been studied extensively. However, it is not clear why depth discrimination (stereoacuity) varies more significantly among people than other modalities. Previous studies reported the involvement of both dorsal and ventral visual areas in stereopsis, implying that not only neural computations in cortical areas but also the anatomical properties of white matter tracts connecting those areas can impact stereopsis and stereoacuity. Here, we studied how human stereoacuity relates to white matter properties by combining psychophysics, diffusion MRI (dMRI), and quantitative MRI (qMRI). We performed a psychophysical experiment to measure stereoacuity, and in the same participants we analyzed the microstructural properties of visual white matter tracts based on two independent measurements, dMRI (fractional anisotropy, FA) and qMRI (macromolecular tissue volume; MTV). Microstructural properties along the right vertical occipital fasciculus (VOF), a major tract connecting dorsal and ventral visual areas, were highly correlated with measures of stereoacuity. This result was consistent for both FA and MTV, suggesting that the structural-behavioral relationship reflects differences in neural tissue density, rather than differences in the morphological configuration of fibers. fMRI confirmed that binocular disparity stimuli activated the dorsal and ventral visual regions near VOF endpoints. No other occipital tracts explained the variance in stereoacuity. In addition, the VOF properties were not associated with differences in performance on a different psychophysical task (contrast detection). These series of experiments suggest that stereoscopic depth discrimination performance is, at least in part, constrained by dorso-ventral communication through the VOF.

## Introduction

Stereopsis is a fundamental human visual function that has been studied over two centuries (1–9). Traditional visual neuroscience has focused on the properties of neural response in gray matter towards important cues for the stereopsis, such as binocular disparity, to understand the neural computation achieving the perception of three-dimensional world (10–14). A series of electrophysiological, functional MRI, and neuropsychological studies have revealed a number of cortical areas involved in binocular disparity processing, which are distributed across dorsal and ventral visual cortices (11–18).

The neural basis of stereopsis is relatively well understood; however, there is one key question that remains unanswered: why the ability to discriminate depth (stereoacuity) varies between people. In fact, a number of psychophysical studies have reported a broad and often bimodal distribution of human stereoacuity, which are much less evident than other visual modalities (19–22). However, the neurobiological origin of such large difference in perceptual performance is unknown.

Given that several visual areas in both dorsal and ventral streams are known to be involved in stereo perception, the anatomical properties of the white matter tract communicating those areas should also be crucial. Recent advancements in non-invasive neuroimaging methods, such as dMRI and qMRI, has opened the avenue to test the relationship between the tissue properties of important white matter tracts and cognitive abilities, because this enables the comparison between neuroanatomical and behavioral measurement in identical human participants (23–28). Here, we have combined modern structural neuroimaging techniques (dMRI and qMRI) with psychophysical measurements to assess human stereoacuity, and clarify how the tissue properties of visual white matter tracts may relate to the stereoacuity (see Materials and Methods). In addition, we have evaluated the relationship between the endpoints of the tracts and areas activated by the same stereo stimuli using fMRI. Furthermore, we tested how these tissue properties relate to the contrast detection threshold to clarify whether the observed relationship between anatomical and psychophysical measurements are specific to stereoacuity.

## Results

### Psychophysical experiment on stereoacuity

We measured the stereoacuity of 19 healthy human participants using a psychophysical experiment in which they judged a perceived depth based on binocular disparity (Fig. 1A). Participants viewed a random dot stereogram or RDS (2) that consisted of a central disk and surrounding ring (Fig. S1A). The surrounding ring was always presented at zero disparity; however, the central disk was presented at a range of disparities across trials. Participants judged the depth of the central disk (“near” or “far”) with respect to the surrounding ring. Stereoacuity was estimated by fitting a psychometric function to each participant’s responses (29). Fig. 1B shows examples of the psychometric functions and estimated stereoacuity discrimination threshold that corresponded to an 84% correct response rate. We confirmed that stereoacuity thresholds varied by more than one order of magnitude across participants (Fig. 1C), consistent with previous psychophysical studies (5, 21, 22). We succeeded in estimating the stereoacuity of 14 participants; five participants had <84% correct responses over the full range of tested disparities (± 7.68 arcmin) and we were unable to determine stereoacuity from their psychometric functions. However, all participants discriminated depth from RDSs with a longer duration (500 ms) and larger disparities (15.36 arcmin) significantly better than chance, indicating that none were stereoblind (see Materials and Methods for Stereoacuity experiment).

**Fig. 1.**
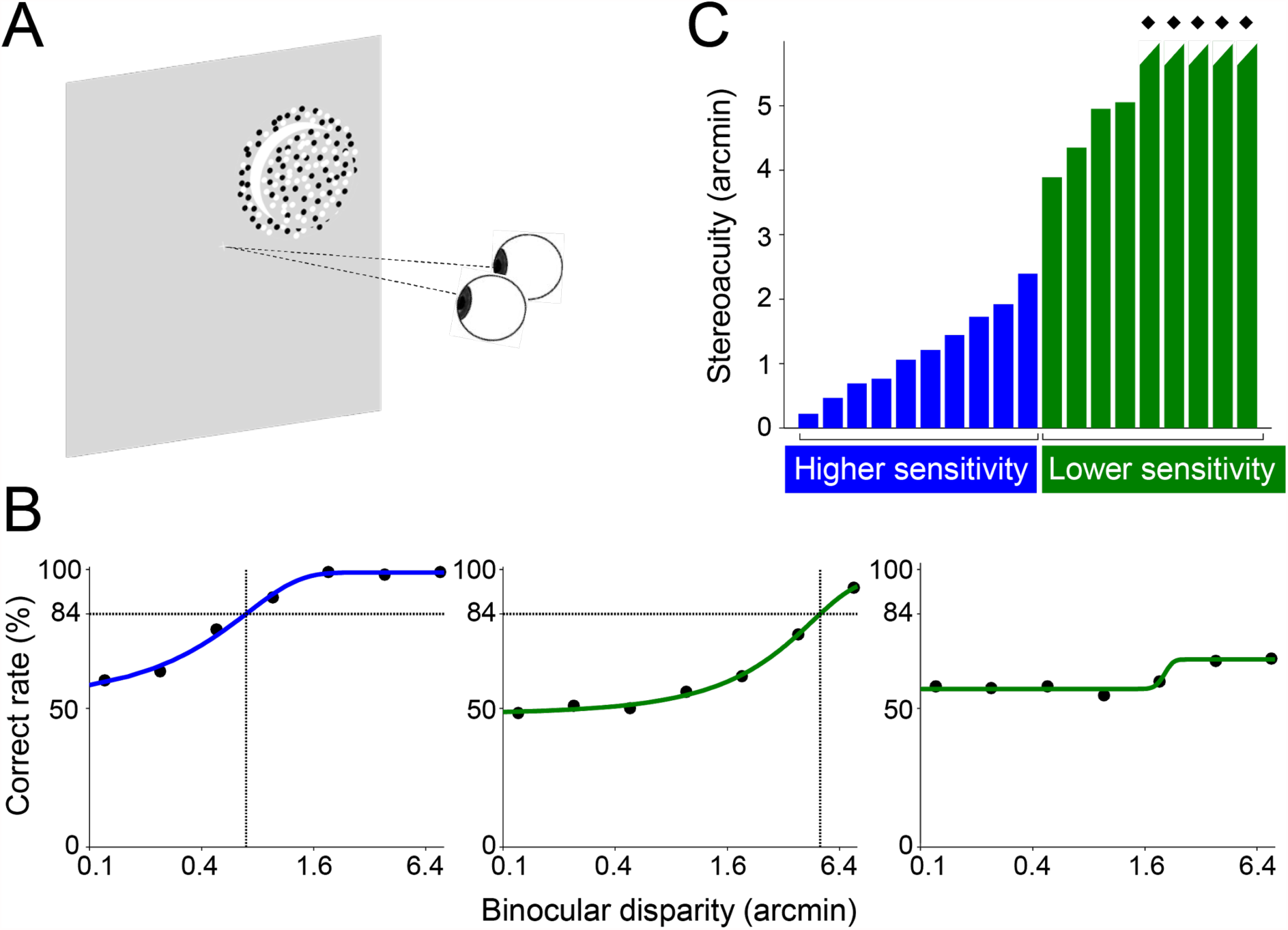
Psychophysical experiment measuring stereoacuity. *(A)* Schematic illustration of the depth discrimination task using RDSs (see Materials and Methods and Fig. S1A for details). Each RDS was concentric-bipartite. Participants were asked to judge whether the central disk was nearer or farther than the surrounding disk. *(B)* The psychometric functions of three representative participants with different performances. The horizontal axis depicts binocular disparity (arcmin; logarithmic scale), while the vertical axis depicts the correct rate. The performance on the crossed and uncrossed disparities was averaged. Stereoacuity was estimated as the binocular disparity at which a participant achieved 84% correct rate. The right panel shows a participant with a performance of <84% over the tested range of disparities; therefore, we could not quantitatively estimate their stereoacuity using the identical criteria. *(C)* The stereoacuity of all participants (*N* = 19). The vertical axis shows the disparity threshold at which performance reached 84% correct. The stereoacuity value is arbitrary for the five participants whose stereoacuity could not be quantitatively estimated (performance <84%; labeled with diamonds). Note, these five participants were not stereoblind (see Materials and Methods for stereoacuity experiment). We classified participants into different subgroups using a two-step clustering algorithm and Schwarz’s Bayesian criterion. The analysis revealed two subgroups, which correspond to good and poor stereoacuity groups, respectively.

### Microstructural properties of VOF explains individual variabilities in stereoacuity

We collected two independent structural MRI datasets, dMRI and qMRI, from the participants who took part in the psychophysical experiment. We performed probabilistic tractography and fascicle evaluations on the dMRI dataset (30, 31) to identify the trajectory of major visual white matter tracts (left and right optic radiation, OR; left and right inferior longitudinal fasciculus, ILF; left and right VOF; and forceps major of the corpus callosum) following the anatomical prescriptions in previous studies (32–37). We evaluated the tissue properties along these visual white matter tracts using the widely used dMRI measure, FA (38), and recently proposed qMRI measure (MTV) that quantifies the non-proton neural tissue density (39, 40). Finally, we examined how the variation of FA or MTV in visual white matter tracts correlates with the stereoacuity thresholds for each participant (see Fig. S2).

First, we examined white matter tracts that explained the variability in stereoacuity by comparing the performance of multiple linear regression models that predict stereoacuity from the tissue properties (MTV or FA) of examined tracts (see Materials and Methods). This analysis was performed on data from the 14 participants whose stereoacuity was quantitatively estimated (an analysis using the data of all 19 participants is also presented below). Next, we selected the best linear regression model using the Bayesian Information Criterion (BIC). BIC model selection for the MTV of visual white matter tracts revealed a significant regression using a single tract, the right VOF (Fig. 2A). This was the best model for predicting the stereoacuity (R^2^ = 0.33, F_(1,12)_ = 5.78, *p* = 0.033; Fig. 2B, Table S1). In addition, BIC model selection revealed that the MTV of the left ILF was a significant predictor of stereoacuity (R^2^ = 0.30, F_(1,12)_ = 5.22, *p* = 0.041; Table S1). No other models using single, or combinations of, visual tracts were significantly correlated with the stereoacuity (Table S1). In summary, the microstructural properties of the right VOF best predicted the variability in stereoacuity.

**Fig. 2.**
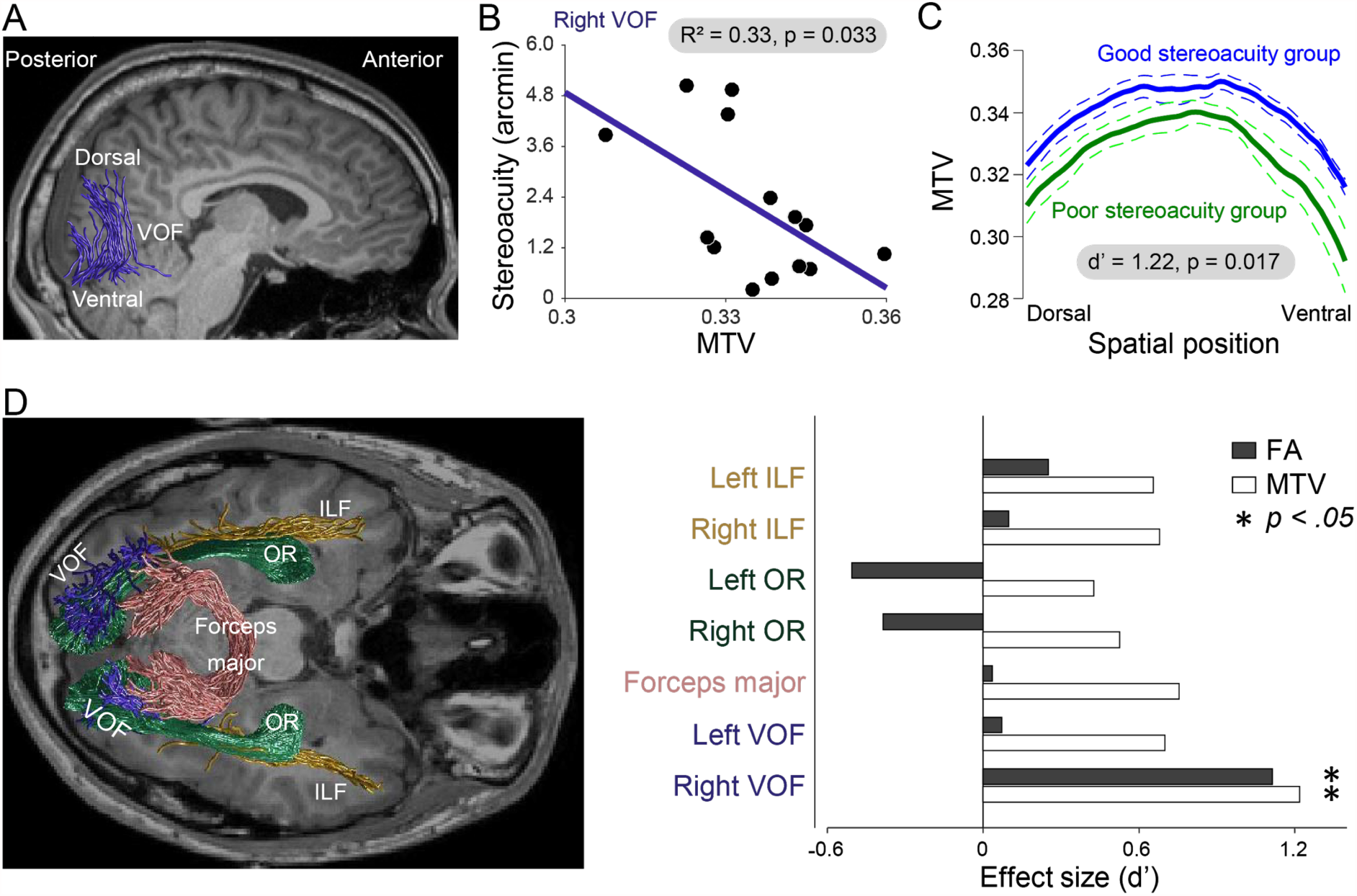
Tissue property of the right VOF explains variabilities in human stereoacuity. *(A)* VOF in the right hemisphere identified in one representative participant (participant 9). VOF connects the dorsal and ventral regions of the occipital cortex. *(B)* Correlation between MTV of right VOF and stereoacuity (*N* = 14; R^2^ = 0.33, *p* = 0.033). See Fig. S3A for the correlation between FA of the right VOF and stereoacuity. *(C)* Tissue properties along the right VOF in good and poor stereoacuity groups. The vertical and horizontal axes represent the MTV and spatial position along the right VOF, respectively. The good stereoacuity group (blue, *n* = 10) showed significantly higher MTV than the poor stereoacuity group (green, *n* = 9) along the entire portion of the right VOF (d’ = 1.22, *p* = 0.017, two-sample *t*-test). Data are represented as mean (solid line) ± S.E.M (dotted line). *(D)* There were no significant stereoacuity-dependent differences in the FA or MTV of other visual white matter tracts. The left panel depicts the visual white matter tracts estimated by tractography (green, OR; pink, forceps major; dark yellow, ILF; blue, VOF) in one representative participant (participant 9). In the right panel, the horizontal axis represents the effect size (d’) of the FA and MTV differences in each visual white matter tract between the good and poor stereoacuity groups. Positive or negative values represent index values that are larger or smaller in the good or poor stereoacuity group, respectively. \**p* = 0.05, two-sample *t*-test. The effect size was largest in the right VOF in a consistent manner across the two independent measurements (FA and MTV).

The BIC model selection using FA, a conventional measure of dMRI, provided similar results: the best model to explain variations in human stereoacuity included a single tract, the right VOF (R^2^ = 0.28, F_(1,12)_ = 4.71, *p* = 0.051; see Fig. S3 and Table S1). No other models, including the model using the FA of the left ILF (R^2^ = 0.12, F_(1,12)_ = 1.57, *p* = 0.23), significantly predicted stereoacuity (Table S1). These results, across two independent measurements using different pulse sequences (dMRI and qMRI), suggested that the observed correlation was related to neural tissue volume along the right VOF, rather than the morphological factors specifically affecting FA (e.g., crossing fibers).

We further examined how MTV and FA differed between the good stereoacuity (low disparity-threshold) and poor stereoacuity (high disparity-threshold) groups, by incorporating datasets from all participants (*N* = 19), including the five participants whose stereoacuity could not be estimated from the psychometric function analysis (Fig. 1B). We classified the participants with quantitative estimates of stereoacuity (*n* = 14) into different subgroups by applying a two-step clustering algorithm to the stereoacuity data and selecting the best clustering based on Schwarz’s Bayesian criterion (see Materials and Methods for details). The analysis revealed two subgroups, which correspond to good (*n* = 10) or poor (*n* = 4) stereoacuity groups. The five participants without quantitative estimates of stereoacuity were included in the poor stereoacuity group (*n* = 9 in total, Fig. 1C). We found that the good stereoacuity group had a significantly higher MTV (Fig. 2C; d’ = 1.22, t_17_ = 2.65, *p* = 0.017) and FA (Fig. S3B; d’ = 1.11; t_17_ = 2.42; *p* = 0.027) along the right VOF when compared with the poor stereoacuity group. This supported our analysis of the relationship between the microstructural properties of the right VOF and stereoacuity (Fig. 2B). The spatial profile of the tract properties suggested that the group difference was present along the entire length of the right VOF, from dorsal to ventral (Fig. 2C and S3B), and not restricted to a localized region. Thus, it is unlikely that the group difference can be explained by a partial volume effect with other short-range fibers (such as U-fibers). We did not find any significant differences in MTV and FA between the two groups in any other visual white matter tracts, such as the left VOF, forceps major, OR, and ILF, in both hemispheres (Fig. 2D). It should be noted that the difference in stereoacuity between these two groups was not accompanied by differences in refractive power or pupillary distance of the eyes (Fig. S1B) nor age (t_17_ = 0, *p* = 1; 25 ± 5.19 and 25 ± 3.71 years old for good and poor stereoacuity groups, respectively).

### VOF connects cortical regions responding to visual stimuli with binocular disparity

To test our hypothesis that the right VOF connects cortical areas that are involved in binocular disparity processing, we performed fMRI experiments to measure the cortical areas activated by the same RDSs as used in the psychophysical experiment (see Materials and Methods for functional MRI data experiment). We observed significant BOLD responses to the RDSs, when compared with uncorrelated RDSs, in both dorsal and ventral extrastriate cortices that were consistent with previous fMRI studies in humans (Fig. 3B) (17, 41–43). Importantly, both dorsal and ventral VOF endpoints overlapped with disparity-selective regions (Fig. 3A-C; see *Supplementary Information*). These results agree with our hypothesis that discrimination of stereoscopic depth involves an interaction between dorsal and ventral cortices through the VOF.

**Fig. 3.**
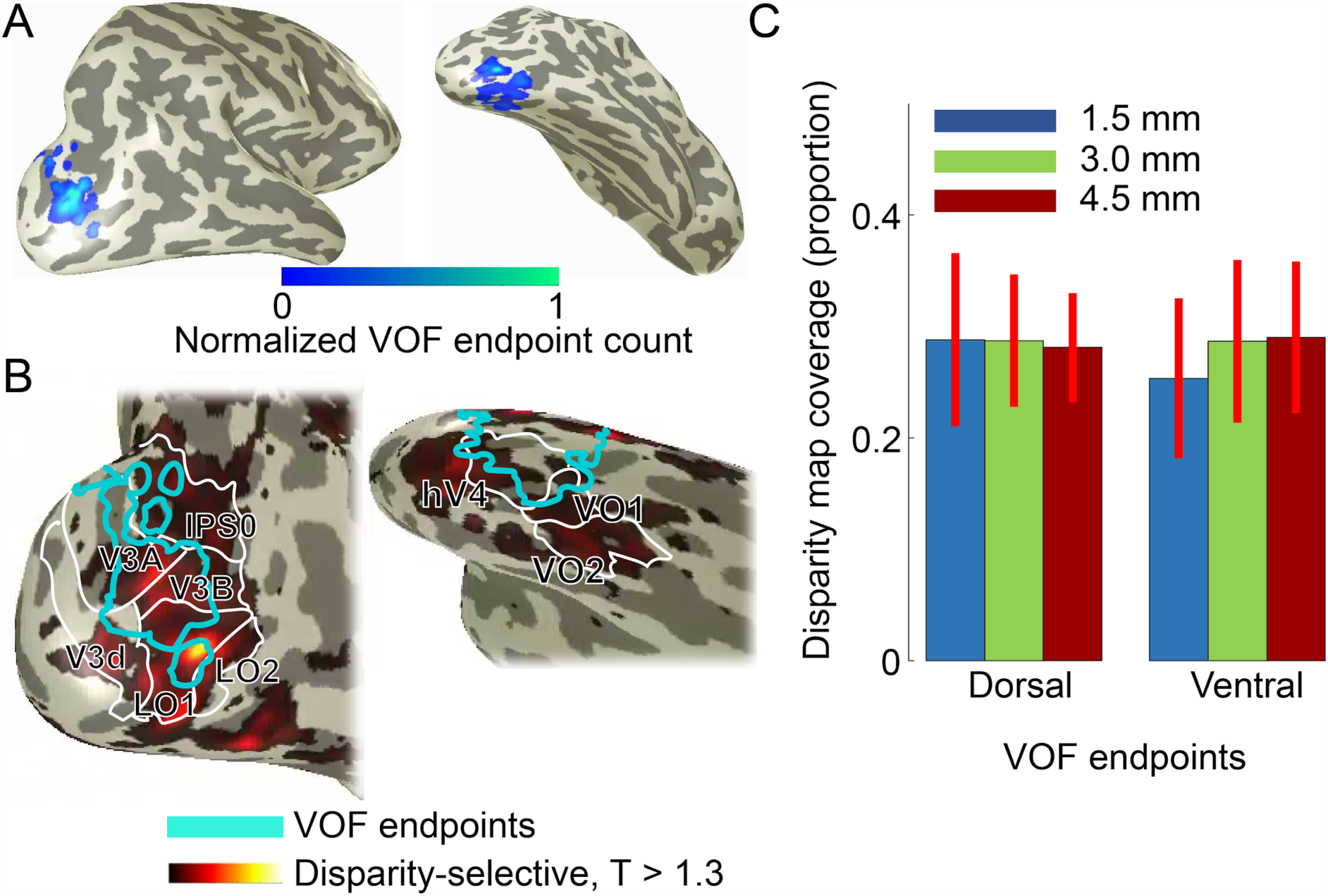
Comparison between VOF endpoints and disparity-sensitive regions. *(A)* VOF endpoints in the right hemisphere from one representative participant (Left panel: dorsal view; Right panel: ventral view; both data from participant 8). Color maps on the cortical surface indicate a normalized count of VOF streamlines having endpoints with 3 mm from each gray matter voxel (see Materials and Methods). The left and right panels represent dorsal and ventral VOF endpoints, respectively. *(B)* Disparity selective areas (hot color) which were significantly activated during the “RDS” blocks when compared with the “uncorrelated RDS” blocks (*p* < 0.05, one-sample *t*-test). The borders of estimated VOF endpoints (cyan, identical in A) as well as the borders of visual areas (white) were overlaid. Cortical coverage of the VOF endpoint and its relation to the retinotopic areas is consistent with a previous study (45). *(C)* Overlap between VOF endpoints and disparity selective regions in the right hemisphere. The vertical axis indicates the proportion of the gray matter voxels near VOF endpoints which intersected with binocular disparity selective areas. The left and right three bars represent dorsal and ventral VOF endpoints, respectively. Different colored bars show data from varying distance thresholds for selecting the gray matter voxels near VOF endpoints (1.5 mm, blue; 3.0 mm, green; 4.5 mm, red). Disparity-selective regions overlap with similar proportion of VOF endpoints across the dorsal and ventral visual cortex. Data represent mean ± S.E.M (*N* = 6).

### Psychophysical experiment on contrast detection sensitivity

Finally, we addressed whether the tissue properties of the right VOF are related specifically to stereoacuity or to a general sensitivity to the visual system. We measured the participants’ contrast detection thresholds using Gabor patch stimuli, which do not require binocular integration (Fig. 4A). In contrast to disparity thresholds, contrast detection thresholds showed a unimodal distribution (Fig. 4B), and were not significantly correlated with stereoacuity (*r* = 0.18, *p* = 0.58). A simple linear model that included the tissue properties of the right VOF did not significantly predict the contrast detection threshold (R^2^ = 0.017, *p* = 0.59 for FA; R^2^ = 0.031, *p* = 0.47 for MTV, Fig. 4C). Group difference analysis revealed no significant difference in the tissue property between the good (low contrast-threshold, *n* = 12) and poor (high contrast-threshold, *n* = 7) contrast sensitivity groups (Fig. 4D; d’ = 0.21, t_17_ = 0.44, *p* = 0.66 for MTV; d’ = 0.16, t_17_ = 0.34, *p* = 0.74 for FA; see Materials and Methods for methods to classify participants into two groups). Taken together, these data show that the variability in FA and MTV values in the right VOF between the good and poor stereoacuity groups (Fig. 2) does not reflect general visual sensitivity.

**Fig. 4.**
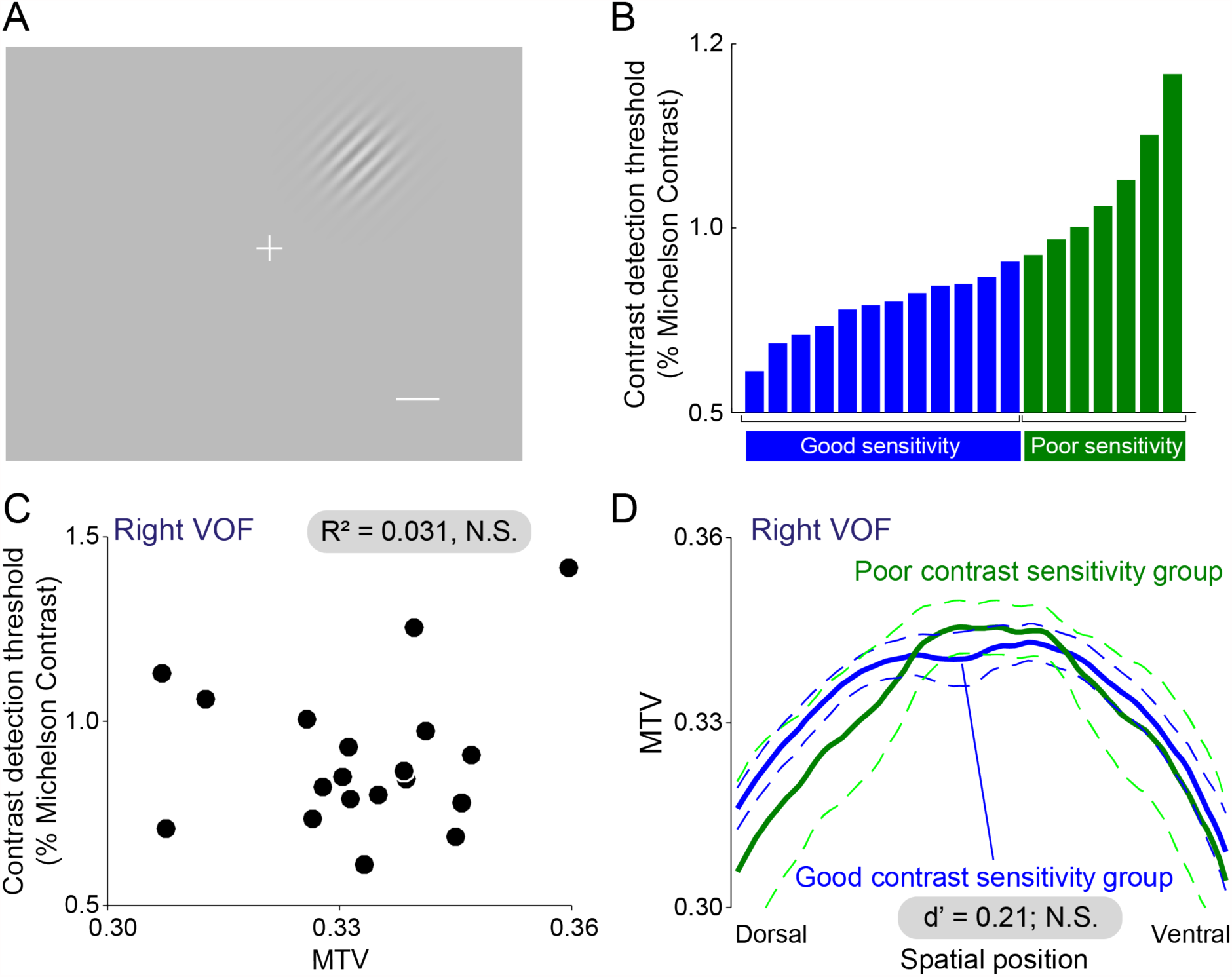
The tissue properties of the right VOF do not explain variabilities in contrast sensitivity. *(A)* Gabor patch stimuli were used to measure contrast sensitivity (see Materials and Methods). The white scale bar depicts 1°, which was not visible in the experiment. *(B)* The contrast sensitivity of all participants (*N* = 19). Using a two-step clustering algorithm and Schwarz’s Bayesian criterion, we classified participants into different subgroups. The analysis indicated two subgroups, which correspond to good and poor contrast sensitivity groups respectively. *(C)* Scatter plot of MTV along the right VOF (horizontal axis) and contrast detection threshold of each participant. The MTV along the right VOF did not significantly predict the contrast detection threshold (R^2^ = 0.031, *p* = 0.47). *(D)* MTV along right VOF in the good and poor contrast sensitivity groups. We did not find a significant difference in MTV between the groups (d’ = 0.21, *p* = 0.66). Data represent mean ± S.E.M. The conventions are identical to those in Fig. 2C.

## Discussion

In the current study, we examined the neurobiological correlates of the large variability in stereoacuity. Advanced non-invasive neuroimaging methods, such as dMRI and qMRI, are advantageous when investigating the neurobiological origin of individual variance in sensory abilities because neuroanatomical and behavioral measurements collected from the same participants can be compared (23–28). We have used this advantage to compare human stereoacuity and white matter properties. We found a significant statistical relationship between stereoacuity and the microstructural properties in the right VOF, a specific white matter tract that connects the dorsal and ventral visual cortex (36, 44, 45). These data support classical and recent theories that emphasize the importance of white matter tracts in understanding sensory and cognitive functions (46, 47). Behavior-anatomy correlates could be found using the MTV and FA, two independent microstructural measurements, in both regression and group comparison analyses. Furthermore, we confirmed that the VOF had endpoints in disparity responsive regions of the dorsal and ventral cortices, suggesting that the VOF connects cortical regions that are involved in disparity processing. Finally, we found that the tissue properties of the right VOF were not related to contrast sensitivity.

A number of previous studies have used conventional diffusion tensor metrics, such as FA, to examine the tissue properties of white matter tracts relative to behavioral characteristics (23, 24, 36, 48). In contrast, few recent studies have used advanced qMRI metrics, such as MTV (39, 40), to assess the microstructural properties of white matter tracts. FA is a reproducible metric with high sensitivity for detecting the tissue structural differences in white matter tracts (49–53). However, the microstructural interpretation of differences in FA is challenging because FA measurements can be associated with many biological factors, such as axon diameter, axon density, myelin-sheath thickness, and tightness of fasciculation due to crossing fibers (38, 51, 54). Here, we combined dMRI with qMRI, which can provide additional information for inferring microstructural properties (39, 55, 56). The MTV is a robust qMRI-based metric that quantifies local tissue volume within each voxel *via* quantification of proton density (39). There is converging evidence indicating that MTV is a reliable approximation of lipid and macromolecular volume fractions (40, 57). Taken together, the relationship between the right VOF and stereoacuity, as shown in both FA and MTV analyses, may reflect a difference in lipid or macromolecule volume fractions, such as myelin thickness or axon density, rather than the morphological configuration of axons, such as the degree of fiber crossings.

Duan and colleagues (58) have investigated the microstructural properties of visual white matter tracts between an amblyopia and control groups. They observed a difference in the diffusion property (mean diffusivity) along the right VOF; however, this difference is not supported by qMRI measurements. Additionally, Duan and colleagues reported a difference in the diffusion property along the optic radiation, which we did not find in this study. The discrepancy between our results and those of Duan and colleagues (58) suggests that the microstructural basis of stereoacuity is distinct from that of amblyopia.

Some visual neuroscience studies have emphasized the role of the “dorsal stream” on stereopsis (59–61), which is consistent with experimental evidence showing the neural correlates of stereoscopic depth perception in dorsal areas (62); however, there are converging lines of evidence showing that not only dorsal but also ventral visual cortices are involved in binocular disparity processing, including computations of relative disparity and disparity-defined three-dimensional shapes (11, 12, 15, 17, 41, 42, 63–70). Moreover, cortical representations that account for perceptually relevant disparity processing are found in higher regions of both the dorsal and ventral visual streams (42, 71), including cortical areas near the VOF endpoints (45). Here, we found that stereoacuity, measured with stimuli of relative disparity (Fig. S1A), is correlated with the properties of the VOF (Fig. 2). In addition, we have confirmed that the endpoints of the VOF overlap with relative disparity selective regions (Fig. 3B and 3C). Therefore, the VOF may transmit perceptually relevant signals of relative disparity by connecting the dorsal and ventral cortices involved in binocular disparity processing.

While we have shown a significant relationship between the right VOF and stereoacuity, we did not find statistically significant results for the left VOF. This asymmetry is consistent with previous studies that report the lateralization of disparity processing to the right hemisphere. For example, neuropsychological studies have reported that brain damage causing the loss of stereoscopic depth perception is lateralized to the right hemisphere (18, 64, 72–77). Furthermore, fMRI studies have demonstrated some evidence for the lateralization of the cortical response to disparity-defined stimuli to the right hemisphere (41, 43, 78–80). Therefore, our results showing that stereoacuity correlates more strongly with the right VOF are in line with these previous publications.

In summary, the anatomical properties of the VOF explain the variability in human stereoacuity thresholds. Stereoacuity may be related to the microstructural properties of anatomical infrastructure for communication between the dorsal and ventral visual cortices.

## Materials and Methods

### Participants

Twenty-three healthy volunteers (19 male, 4 female; mean age, 26.1 years) participated in the study. None of the participants had a history of eye disease. All participants gave written informed consent to take part in this study, which was conducted in accordance with the ethical standards stated in the Declaration of Helsinki and approved by the local ethics and safety committees at the Center for Information and Neural Networks (CiNet), National Institute of Information and Communications Technology.

### Stereoacuity experiment

Nineteen participants (16 males, 3 females; mean age, 25.0 years) took part in the experiment to determine their stereoacuity. All procedures were completed in approximately 80 minutes for each participant. The stereoacuity experiment employed a haploscope: each eye viewed one half of the monitor through an angled mirror and a front triangular prism mirror. Each RDS was composed of a central disk (diameter: 3°) and surrounding ring (width: 0.5°, outer diameter: 4°; see Fig. S1A for an example). The surrounding ring always had zero disparity. The binocular disparity in the central disk varied across trials (disparity magnitudes: ±0.12, ±0.24, ±0.48, ±0.96, ±1.92, ±3.84, and ±7.68 arcmin), which were chosen based on a typical range of human stereoacuity (21). Participants performed a single-interval, two-alternative forced-choice near/far discrimination task. Participants judged whether the central stimulus appeared nearer or further than the surrounding ring while fixating on the central fixation point (Fig. S1A). The stimulus position (Up-Right, Up-Left, Down-Right and Down-Left) was constant within a session of 42 trials (three trials for each disparity condition) and was randomized across sessions. Participants completed 20 sessions (5 sessions for each stimulus position). We estimated stereoacuity in each participant by fitting their correct response rate to a cumulative Gaussian psychometric function using a maximum-likelihood estimation (29). The responses were pooled across all four positions and crossed and uncrossed disparities because there was no systematic difference in performance across positions or depth directions. Hereafter, we defined stereoacuity as the magnitude of binocular disparity that corresponded to the 84% correct rate in the task. We identified stereoacuity for 14 participants using this procedure, however, we could not identify stereoacuity in the remaining 5 participants because their correct response rate was <84% in all disparity conditions (Fig. 1). We confirmed that the five participants were not stereoblind because they could accurately discriminate depth with a longer duration (500 ms) and larger disparity presented in a practice session. Correct rate at 15.36 arcmin was 100%, 70.8%, 83.3%, 87.5%, and 95.8% for those each participant which were significantly higher than chance (95% confidence interval = 42.7–63.3% using a binomial distribution). See *Supplementary Information* for further technical details.

### Contrast threshold experiment

Nineteen participants (15 males, 4 females; mean age, 26.0 years old) underwent a contrast threshold experiment. Fifteen of these participants also participated in the stereoacuity experiment. All procedures were completed in approximately 57 minutes for each participant. We presented Gabor patch stimuli whose orientation was tilted 45° to the left or right from vertical (Fig. 4A). The stimulus positions were identical to those used in the stereoacuity experiment. The experiment composed of two stages. An approximate threshold was measured in the first stage, which was used to determine the contrast range that was used at the second stage to estimate a precise threshold. In both stages, participants were asked to judge whether the stimulus orientation was tilted towards the left or right. The details in apparatus, stimulus and two-step procedure for threshold estimation is described in *Supplementary Information*.

### Structural MRI experiment

All MRI data were acquired using a 3T SIEMENS Trio Tim scanner at the Center for Information and Neural Networks (CiNet), National Institute of Information and Communications Technology, and Osaka University.

#### Anatomical MRI data acquisition and tissue segmentation

We collected T1-weighted MR-RAGE image (1 mm isotropic; TR, 1900 ms; TE, 2.48 ms) from all participants (*N* = 23) to estimate white/gray matter border. The segmentation was performed using an automated procedure in Freesurfer software (https://surfer.nmr.mgh.harvard.edu/) (81). The tissue segmentation was used for subsequent dMRI and fMRI (see *Supplementary Information*) analyses. Acquisition of the anatomical MRI data took approximately 15 min per participant.

#### Diffusion MRI data acquisition

We collected dMRI data from all participants (*N* = 23) using a 32-channel head coil. Data were acquired using dual-spin echo planar imaging (EPI; TR, 5000 ms; TE, 73 ms; multi-band factor, 2; partial Fourier, 5/8; voxel size, 2 × 2 × 2 mm^3^) implemented in multi-band accelerated EPI pulse sequence provided by the Center for Magnetic Resonance Research, Department of Radiology, University of Minnesota (https://www.cmrr.umn.edu/multiband/) (82). The diffusion weighting was isotropically distributed along the 64 directions (b = 1000 s/mm^2^). Non diffusion-weighted (b = 0) images were acquired at the beginning and end of the dMRI session (two b = 0 volumes per image set). To minimize EPI distortion, two image sets were acquired with reversed phase-encoding directions (A-P and P-A). Acquisition of the dMRI data took approximately 20 min for each participant.

#### Quantitative MRI data acquisition

We collected qMRI data from all participants (*N* = 23) using a 32-channel head coil. QMRI measurements were obtained using protocols described in a previous publication (39). We measured four Fast Low Angle Shot (FLASH) images with flip angles of 4°, 10°, 20°, and 30° (TR, 12 ms; TE, 2.41 ms), and a scan resolution of 1 mm isotropic. We collected five additional spin echo inversion recovery (SEIR) scans with an echo planar imaging (EPI) readout (TR, 3 s; TE, 49 ms; 2× acceleration) to remove field inhomogeneities. The inversion times were 50, 200, 400, 1200, and 2400 ms. In-plane resolution and slice thickness of the additional scan was 2 × 2 mm^2^ and 4 mm, respectively. Acquisition of the qMRI data took approximately 35 min for each participant.

### Diffusion MRI data analysis

DMRI data preprocessing was performed using mrDiffusion tools implemented in the vistasoft distribution (https://github.com/vistalab/vistasoft). We identified visual white matter tracts in each participant, from whole-brain streamline generated by probabilistic tractography implemented in MRtrix3 (http://www.mrtrix.org/) and selected by Linear Fascicle Evaluation (LiFE; https://francopestilli.github.io/life/) (31). Details are described in *Supplementary Information*.

### Quantitative MRI data analysis

QMRI data were processed using the mrQ software package (https://github.com/mezera/mrQ) produce the macromolecular tissue volume (MTV) maps (39). Details are described in *Supplementary Information*.

### Functional MRI experiment

We collected fMRI data from eight participants who participated in the stereoacuity psychophysical experiment (7 male, 1 female; mean age, 26.6 years old). We used an identical scanner to the structural MRI experiment with the posterior section of a 32-channel coil. Data were acquired at a resolution of 2.0 mm isotropic voxels with an interleaved T2*-weighted gradient echo sequence. Participants viewed gray background, RDS, or uncorrelated RDS (uRDS) through a polarized 3D system, during which they performed a fixation task requiring vernier detection (42). We excluded two participants with poor task performance during the fMRI scan (<75% correct rate) from subsequent analyses. The disparity-selective areas are defined as cortical gray matter voxels that responded more strongly to RDS than uRDS (*p* < 0.05, one-sample *t*-test). Acquisition of fMRI data took approximately 60 min for each participant. See *Supplementary Information* for further details.

### Evaluating the tissue properties of white matter tracts

We evaluated the tissue properties of each visual white matter tract based on methods used in previous studies (35, 58). Briefly, we resampled each streamline to 100 equidistant nodes. Tissue properties were calculated at each node of each streamline using spline interpolation of the tissue properties: FA and MTV. MTV maps were registered with dMRI data for each participant and we computed the MTV values along each node of each streamline. The properties at each node were summarized by taking a weighted average of the FA or MTV on each streamline within that node. The weight of each streamline was based on the Mahalanobis distance from the tract core. We excluded the first and last 10 nodes from the tissue property of tract core to exclude voxels close to gray/white matter interface where the tract is likely to be heavily intersected with the superficial U-fiber system. We summarized the profile of each tract with a vector of 80 values representing the FA or MTV values sampled at equidistant locations along the central portion of the tract. FA was averaged across two sessions.

### Regression analysis

First, we examined whether differences in white matter tracts could explain the variability in stereoacuity between participants using multiple linear regression models, which predicted stereoacuity from tract properties. In this analysis, we used the data from the 14 participants whose stereoacuity had been successfully estimated. For each participant, MTV or FA values along 80 equidistant nodes in each tract were averaged to obtain a participant-specific single number summary representing tract tissue property. Following this, we generated multiple linear regression models that could predict the variability in stereoacuity by using the tract properties as explanatory variables. In each model, explanatory variables were chosen from the tract property (either MTV or FA) of the individual, or combinations of the white matter tracts we identified (left and right OR, left and right ILF, left and right VOF, and forceps major). In total, we tested 127 possible linear regression models for each MTV and FA to predict stereoacuity. We used the Bayesian Information Criterion (BIC), which enables a model comparison by penalizing models with more parameters as a redundant model, to select the best model. We define the model with the smallest BIC as the best model for MTV and FA separately. In addition, we evaluated the statistical significance of the goodness-of-fit of each model using the F-test (Table S1). We also performed a regression analysis for contrast sensitivity data using identical procedures.

### Group comparison analysis

#### Grouping of participants based on stereoacuity

In this analysis, we utilized data from all 19 participants, including the 5 participants whose stereoacuity could not be quantitatively estimated. First, we classified the 14 participants into different subgroups by applying a two-step clustering algorithm to stereoacuity data and selecting the best clustering based on Schwarz’s Bayesian criterion (SPSS Statistics 25, IBM, USA) (83). This method revealed two subgroups, which corresponded to good (*n* = 10; 1.19 ± 0.69 arcmin) and poor (*n* = 4; 4.56 ± 0.55 arcmin) stereoacuity. Next, we classified the five participants without quantitative stereoacuity estimates into the poor stereoacuity group (*n* = 9; Fig. 1C).

#### Grouping of participants in contrast threshold experiment

We classified participants into the different subgroups by applying a two-step clustering algorithm to the contrast threshold data and selecting the best clustering based on Schwarz’s Bayesian criterion. This method revealed two subgroups, corresponding to good (*n* = 12; 0.78% ± 0.085 Michelson contrast) and poor (*n* = 7; 1.11% ± 0.17 Michelson contrast) contrast sensitivity, respectively (Fig. 4B).

#### Statistical comparisons

We compared the tract profiles (FA or MTV) between the good and poor performance groups, as defined in the psychophysical experiments (stereoacuity or contrast sensitivity; see above for grouping of participants). We calculated the mean FA and MTV across all 80 nodes for each participant and tract. We computed the effect size (d’) and statistical significance of the inter-group differences using a two-sample *t*-test.

## Acknowledgements

This study was supported by Japan Society for the Promotion of Science (JSPS) KAKENHI (JP17H04684 to HT, JP16H01673 and JP17H01381 to IF, JP16H05862 to KA), a grant from the Ministry of Internal Affairs and Communications (to IF), and Grant-in-Aid for JSPS Fellows (JP15J00412 to HT). We thank Atsushi Wada for providing the computing environment to run the analyses, and thank Aviv Mezer, Garikoitz Lerma-Usabiaga and Takashi Ueguchi for the suggestion on quantitative MRI data collection, and Thomas Baumgartner for his participation in an initial stage of this study. We also thank Hiroshi Ban, Nobuhiro Hagura, Franco Pestilli, and Brian A. Wandell for comments on an earlier version of the manuscript. This research was supported in part by “Program for Leading Graduate Schools” of the Ministry of Education, Culture, Science and Technology, Japan (to HO).

## Notes

**Conflict of interest** The authors declare no conflict of interest.

