## Supplementary Materials for "Microstructural properties of the vertical occipital fasciculus explain the variability in human stereoacuity"


・ **Author Information**

Hiroki Oishi[a,b,1)], Hiromasa Takemura[a,b,1)], Shuntaro C. Aoki[a)], Ichiro Fujita[a)], Kaoru Amano[a,b)]

[a)] Graduate School of Frontier Biosciences, Osaka University, Suita 565-0871, Japan

[b)] Center for Information and Neural Network (CiNet), National Institute of Information and Communications Technology, and Osaka University, Suita 565-0871, Japan

[1)] These authors contributed equally

Supplementary Materials and Methods

***Stereoacuity experiment***

*Apparatus*

RDSs were presented with a spatial resolution of 3840 × 2160 pixels and frame rate of 60 Hz on a gamma-corrected, full-flat 4K LCD monitor (P2715Q, Dell, USA). The distance from the angled mirror to the prism mirror was 10 cm. The direct distance from the display to chin support was 160 cm; therefore, the viewing distance was 170 cm. Each side of the monitor covered a visual angle of 10.0° × 11.3°. RDSs were presented at one of 4 different positions (Up-Right, Up-Left, Down-Right, and Down-Left), whose center was 3° away from the fixation point (see Fig. S1A).

*Stimuli*

The RDS stimulus consisted of an equal number of black (0.15 $cd \cdot m^{-2}$) and white (171.85 $cd \cdot m^{-2}$) dots on a mid-gray background (86.00 $cd \cdot m^{-2}$). The diameter and density of the dots was 0.16° and 25%, respectively. Dot patterns in each eye were refreshed at 20 Hz. The stimulus duration was 94 ms, which avoided the occurrence of vergence eye movements (84). The inter-stimulus interval was 2 s.

*Practice task*

Prior to the main experiment, the participants practiced the task with a longer stimulus presentation time (500 ms) and 8 different binocular disparities (±0.24, ±0.96, ±3.84, and ±15.36 arcmin). Every participant practiced for an identical number of trials (24 trials, including 3 trials for each disparity condition).

***Contrast threshold experiment***

*Apparatus*

We used a gamma-corrected (https://github.com/hiroshiban/Mcalibrator2) (85), full-flat 10-bit LCD monitor (sx2262w, Eizo, Japan; spatial resolution, 1920 × 1200 pixels, which corresponds to 43.4° × 27.4°; frame rate, 60 Hz) to present Gabor patch stimuli. The viewing distance was 60 cm.

*Stimuli*

We presented Gabor patch stimuli whose orientation was tilted 45° to the left or right from vertical (Fig. 4A; spatial frequency, 3 cycles/°; stimulus diameter, 4°; the standard deviation of Gabor filter, 0.8°). The mean luminance of the Gabor patch stimuli and background was 112.37 $cd \cdot m^{-2}$. The stimulus duration was 100 ms.

*Task design and estimation of threshold*

In the first stage of the experiment, we presented Gabor stimuli of six different levels of luminance contrast (0.00%, 0.39%, 0.77%, 1.16%, 1.54%, and 1.91% Michelson contrast). One session of 18 trials (3 trials for each contrast condition) was performed for each stimulus position. We pooled data from the four different positions and approximated the threshold as the contrast corresponding to the 84% correct rate by fitting to a psychometric function (29).

In the second stage of the experiment, the stimulus contrast was varied at 12 levels between 0–1.73% (12 participants), 0–2.1% (6 participants), or 0–2.47% (6 participants) to match the approximate threshold identified in the first stage. We estimated the contrast detection threshold (84% correct response rate) by fitting a cumulative Gaussian function to the correct response rate obtained at the second stage.

### *Diffusion MRI data analysis*

Preprocessing

DMRI images were corrected for susceptibility-induced distortions using FSL TOPUP tools (86). Eddy current distortions and participant motion in the dMRI images were removed by a 14-parameter constrained non-linear co-registration based on the expected pattern of eddy-current distortions given the phase-encode direction of the acquired data (87) using mrDiffusion tools implemented in the vistasoft distribution (https://github.com/vistalab/vistasoft).

Tracking

We used constrained spherical deconvolution (CSD; $L_{max}$ = 8) (88) to estimate the fiber orientation distribution in each voxel using MRtrix3 (30) (http://www.mrtrix.org/). We performed probabilistic tractography implemented in MRtrix3 to generate 2 million candidate streamlines for each dMRI dataset (step size, 0.2 mm; maximum angle between successive steps, 9°; minimum length, 10 mm; maximum length, 250 mm; FOD amplitude stopping criterion, 0.1). The seed voxels for tracking were randomly chosen from the gray-white matter interface region (89).

Tractography optimization

We optimized the estimate of tractography using a Linear Fascicle Evaluation (LiFE; https://francopestilli.github.io/life/) (31, 90). Briefly, using LiFE, we eliminated streamlines that made no contribution to predict the diffusion signal. Further technical details of LiFE are described in previous publications (31, 45, 91, 92).

### *Quantitative MRI data analysis*

Both the FLASH and SEIR scans were processed using the mrQ software package (https://github.com/mezera/mrQ) in MATLAB to produce the macromolecular tissue volume (MTV) maps (39). The mrQ analysis pipeline corrects for RF coil bias using SEIR-EPI scans, producing accurate proton density (PD) and T1 fits across the brain. We used voxels from within the ventricles to denote cerebrospinal fluid (CSF). MTV maps were produced by calculating the fraction of a voxel that is non-water (CSF voxels were classed as approximately 100% water). The full analysis pipeline can be found at previous publications (39, 93).

### *Tract identification from diffusion MRI data*

We identified major visual white matter tracts from streamlines generated by probabilistic tractography in MRtrix3 and selected by LiFE, except for the optic radiation (see below).

#### Vertical occipital fasciculus

We identified the vertical occipital fasciculus (VOF) using open-source MATLAB code distributed with the Automated Fiber Quantification (AFQ) toolbox (35) (https://github.com/yeatmanlab/AFQ). We identified streamlines that traveled vertically, were located posterior to the arcuate fasciculus, and whose ventral endpoints were near the ventral and lateral occipito-temporal cortices, as defined in the Freesurfer atlas (94). Further details of the VOF identification method are described in previous papers (36, 37, 58).

#### Forceps major

We identified the forceps major as streamlines that passed through three ROIs: the mid-sagittal plane of the corpus callosum (95), which was automatically segmented using mrDiffusion tools; and two ROIs in the left and right hemisphere, which were manually defined on the coronal plane as located at Y = –65 (ACPC coordinate). These ROIs were manually defined because the ROIs transformed from the MNI152 template was located too posterior to identify forceps major in some participants, most likely due to morphological differences between Japanese brains and the MNI152 template.

#### Inferior longitudinal fasciculus

We identified the Inferior longitudinal fasciculus (ILF) using automated pipelines implemented in the AFQ toolbox (https://github.com/yeatmanlab/AFQ) (35). Briefly, AFQ transforms two coronal ROIs (anterior and posterior) from the MNI152 template into individual brains and then selects streamlines passing through these ROIs (35).

#### Optic radiation

We identified the optic radiation (OR) using a dedicated method (ConTrack; (96)), because there are known challenges to estimating the human OR using standard whole-brain tractography, in particular, tracking the crossing fiber regions around Meyer's loop (97). First, we estimated the approximate location of the lateral geniculate nucleus (LGN) *via* manual inspection of the T1-weighted image and deterministic tractography from the optic chiasm (98). Following this, we placed a sphere (radius, 8 mm) over the LGN endpoints of streamlines from the optic chiasm. Second, we identified the location of the primary visual cortex (V1) using a probabilistic atlas of retinotopic visual areas (99). Using ConTrack, we sampled 100,000 candidate streamlines connecting the LGN and V1 (angle threshold, 90°; step size, 1 mm). Tracking was restricted using a white matter mask generated by tissue segmentation. We selected 50,000 streamlines with the highest score in the ConTrack scoring process (34). Further details on the methods to identify the OR using ConTrack are described in previous papers (34, 37, 58, 98, 100).

*Across-session averaging and outlier exclusion*
We identified the tracts (VOF, Forceps Major, ILF, and OR) of each participant separately for two dMRI sessions with reversed phase encoding directions. After merging the streamlines of each tract from the two sessions, we excluded outlier streamlines based on criteria used in previous studies (37, 45) for subsequent evaluation of tissue properties.

### Functional MRI data acquisition

We collected functional MRI (fMRI) data to identify the cortical areas that were selectively activated by visual stimuli with binocular disparity.

*fMRI Acquisition protocol*
fMRI data were acquired using a 3T Siemens Trio scanner and the posterior section of a 32-channel coil. Functional data were collected using the parallel acceleration technique (iPAT; acceleration factor = 2) and a simultaneous multi-slice EPI sequence (multi-band factor = 2) to acquire whole-brain (60 slices) volumes (TR, 2 s; TE, 30 ms; flip angle, 70°; FOV, 192 $mm^2$; acquisition matrix, 96 × 96; slice thickness, 2 mm with no gap). These data were obtained using a multi-band accelerated EPI pulse sequence provided by the Center for Magnetic Resonance Research, Department of Radiology, University of Minnesota (https://www.cmrr.umn.edu/multiband/) (101). The slices were aligned parallel to the AC-PC (anterior commissure-posterior commissure) line.

*Apparatus*
Stimulus presentation inside the scanner *was via* a stereoscopic projector system. Briefly, the left and right images were generated by a dual head graphic converter (Matrox DualHead2Go, Matrox) and were projected *via* polarizing filters on a screen placed in front of the participants. The participants viewed the screen through eye glasses with a pair of polarizing filters that match those on the projector. Stimuli were presented at a spatial resolution and frame rate of 1024 × 820 pixels and 85 Hz, respectively, on a full-flat screen (333 × 266 mm). The viewing distance and visual angle of the screen was 97 cm and 19.5° × 15.6°, respectively.

*Experimental design and task*
The participants viewed 12 s blocks of gray background ("Blank"), RDS, or uncorrelated RDS (uRDS), during which they performed a fixation task requiring vernier detection (see below). In "RDS" or "uRDS" blocks, the same stimuli were presented at all four positions used in the psychophysical experiment. The stimuli were presented 12 times for 0.5 s with an inter-stimulus-interval of 0.5 s. During "RDS" blocks, the RDSs that were identical to those presented in the stereoacuity experiment were presented. Within each block, binocular disparity in central disks was randomly chosen from ±1.92, ±3.84, or ±7.68 arcmin (each

binocular disparity was presented twice). During “uRDS” blocks, the random dot stimuli presented in the left and right eyes were mutually independent so that participants could not perceive depth. During “Blank” blocks, only a fixation point was presented. Each session started with a “Blank” block, followed by 6 repetitions of 3 types of blocks (“RDS” - “Blank” - “uRDS” - “Blank”). Ten sessions were repeated for each participant.

During the fMRI experiment, the participants were asked to perform a demanding vernier detection task (42) on the fixation point. This ensured proper fusion of the left and right images and similar levels of attention engagement across different stimuli (correlated, uncorrelated, and blank) (42).

***Functional MRI data analysis***

Functional MRI data were analyzed using MrVista in vistasoft distribution (https://github.com/vistalab/vistasoft). We corrected the slice timing to match the multi-slice acquisition order and corrected for motion within and between scans. We fitted a general linear model (GLM) consisting of predictors convolved with hemodynamic response function (two-gamma HRF) (102) to the time course of each voxel.

We compared the beta weights of the predictors between the ”RDS” and “uRDS” blocks and generated statistical maps of contrasts ($p < 0.05$, one-sample $t$-test). We used a probabilistic atlas, as proposed in a previous study (99), to estimate the cortical location of visual field maps.

***Evaluating the overlap between VOF cortical endpoints and fMRI-based activation map***

We evaluated the overlap between the fMRI activation map, driven by binocular disparity (see Functional MRI data analysis), and VOF endpoints. First, we measured the distance between VOF streamline endpoints and gray matter voxels. We defined gray matter voxels within 1.5 mm, 3.0 mm, 4.5 mm, of VOF streamline points, respectively, as voxels that were covered by the VOF. Second, we calculated the proportion of gray matter voxels near VOF intersects with a fMRI activation map at a specific statistical threshold ($p < 0.05$; see above). We computed the proportion separately for dorsal and ventral VOF endpoints. This overlap analysis was used in several previous publications (37, 45, 92).

We note that there is a known limitation in the spatial precision of this analysis originating from the challenges found in associating the cortical surface and dMRI-based estimates of tract endpoints (103). This analysis measures the general proximity between cortical maps and tract endpoints; however, it is not a definitive estimate of fiber projections into cortical gray matter regions.

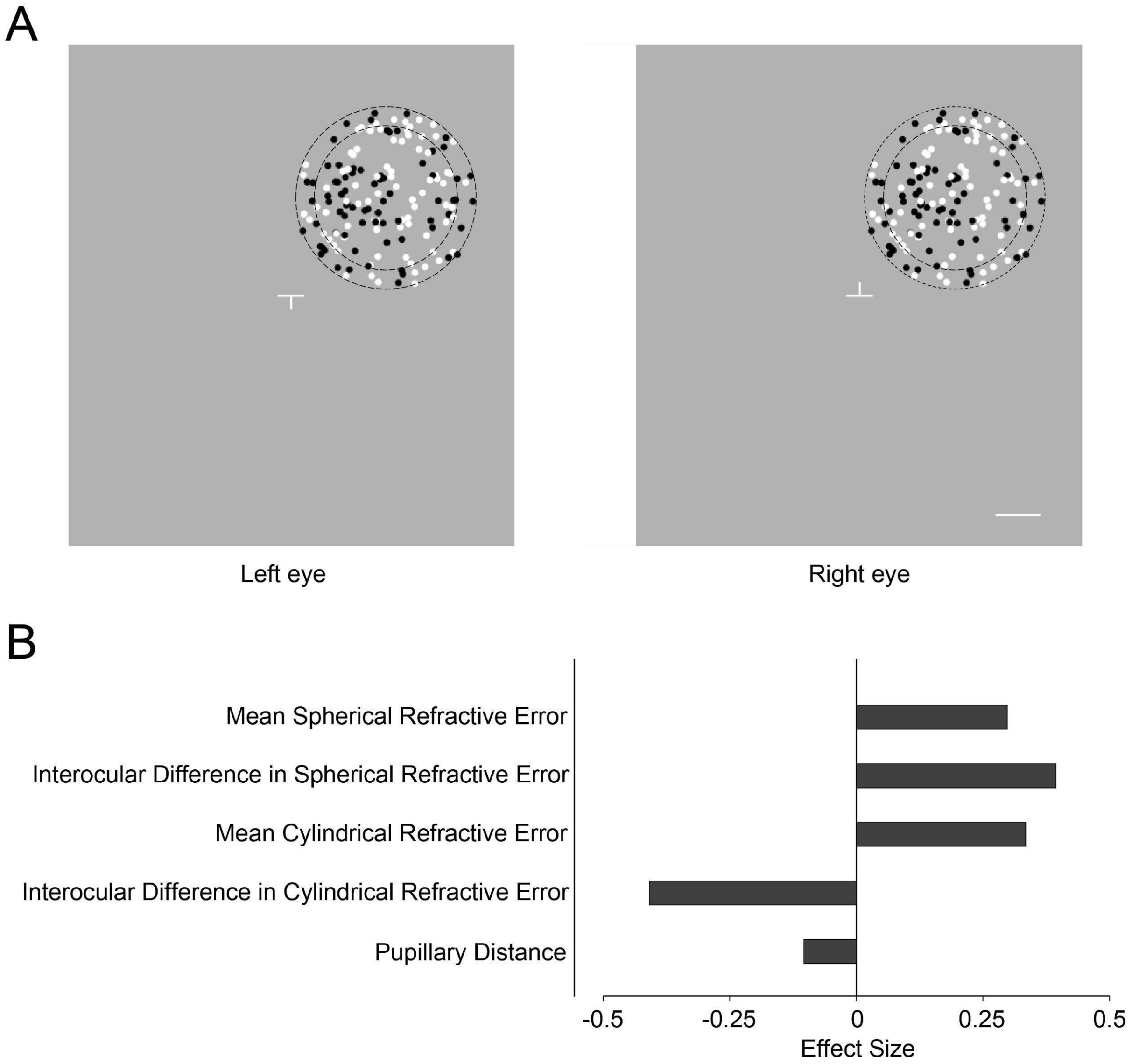


**Fig. S1. Details of stereoacuity experiment.** *(A)* Random dot stereogram (RDS) stimuli used to test stereoacuity. Left and right panels show the stimulus in the left and right eyes, respectively. We used a haploscope to present the RDS stimulus. The RDSs consisted of central and surrounding stimuli, which are highlighted as dotted contours (not visible in the experiment). Dots on the central disk had crossed or uncrossed binocular disparities, while dots on the surround ring and fixation target (a cross when properly fused) had zero disparity (See Materials and Methods for more details of stimulus parameters). The white scale bar depicts 1°, which was not visible in the experiment. Participants were asked to judge whether the central disk appeared to be nearer or farther than the surrounding ring. *(B)* The difference in refractive errors and pupillary distance of the eyes between good (low disparity-threshold) and poor (high disparity-threshold) stereoacuity groups. The difference in stereoacuity between these two groups was not accompanied by a difference in mean spherical refractive error (d’ = 0.30, $t_{17}$ = 0.65, $p$ = 0.53, –4.25 ± 2.16 and –4.93 ± 2.41 diopters for good and poor stereoacuity groups, respectively), mean interocular difference in spherical refractive error (d’ = 0.39, $t_{17}$ = 0.86, $p$ = 0.40, 0.90 ± 0.53 and 0.69 ± 0.51 diopters for good and poor stereoacuity group, respectively), mean cylindrical refractive error (d’ = 0.33, $t_{17}$ = 0.73, $p$ = 0.48, –0.85 ± SD = 0.74 and –1.07 ± 0.59 diopters for good and poor stereoacuity groups, respectively), mean interocular difference in cylindrical refractive error (d’ = –0.41, $t_{17}$ = –0.89, $p = 0.39$, 0.40 ± 0.38 and 0.64 ± 0.75 diopters for good and poor stereoacuity groups, respectively), or mean pupillary distance between eyes (d’ = –0.10, $t_{17}$ = –0.23, $p = 0.82$, 63.3 ± 2.06 and 63.56 ± 2.83 mm for good and poor stereoacuity groups, respectively).

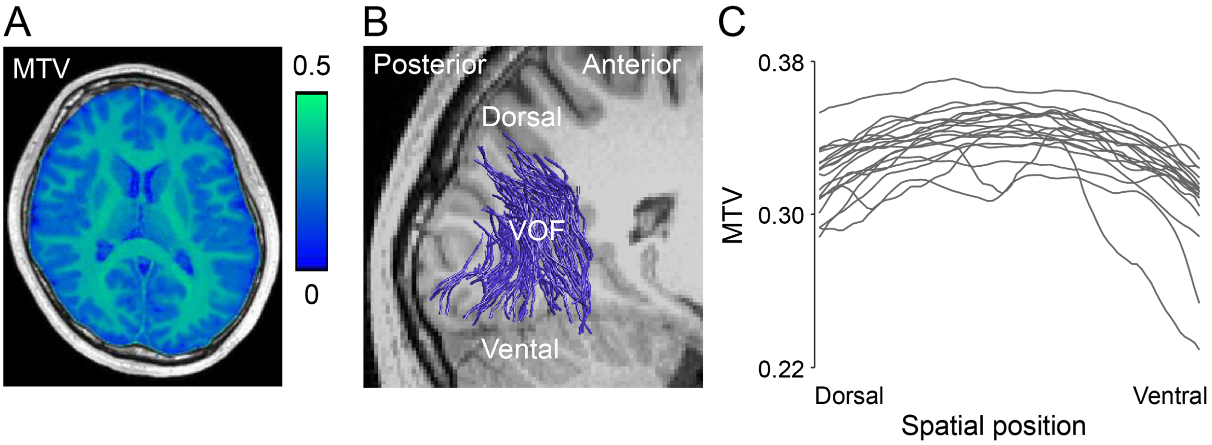


**Fig. S2. Macromolecular tissue volume (MTV) along a tract of interest.** *(A)* A representative MTV values map in the axial plane from a single participant (participant 4). *(B)* The VOF identified from dMRI data in the same participant. *(C)* MTV values of all participants ($N$ = 19) who participated in the stereoacuity experiment. We divided the VOF in each participant into 100 nodes from dorsal to ventral. We estimated the profile of MTV along the VOF by coregistration across the dMRI dataset and MTV maps in each individual participant and calculated the MTV values in each node along the VOF. MTV values (vertical axis) from node 11 (dorsal) to 90 (ventral; horizontal axis) were plotted. Each line indicates the MTV profile an individual participant. See the Materials and Method for technical details regarding the estimation of the MTV profile along white matter tracts.

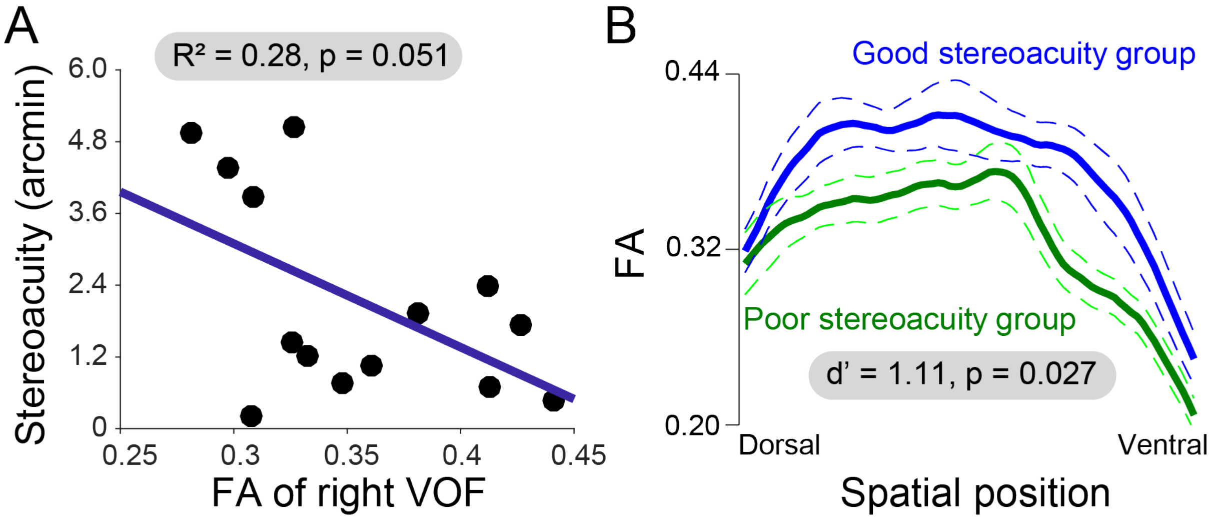


**Fig. S3. Fractional Anisotropy (FA) of the right VOF and variabilities in stereoacuity.** *(A)* There was a correlation between the FA of the right VOF and stereoacuity, but it did not reach statistical significance ($N$ = 14; $R^2$ = 0.28, $p$ = 0.051). *(B)* The FA along the right VOF in the good (blue, $n$ = 10) and poor (green, $n$ = 9) stereoacuity groups. The conventions are identical to Fig. 2.

| | Model fitting | | Model selection |
|---|---|---|---|
| Variables in the model | $R^2$ | *p* (using F-test) | BIC |
| FA | | | |
| Left ILF | 0.12 | 0.23 | 13.79 |
| Right ILF | $0.70 * 10^{-5}$ | 0.98 | 13.91 |
| Left OR | 0.031 | 0.54 | 13.88 |
| Right OR | 0.024 | 0.60 | 13.89 |
| Forceps major | 0.033 | 0.53 | 13.88 |
| Left VOF | $0.23 * 10^{-2}$ | 0.87 | 13.91 |
| Right VOF | 0.28 | 0.051 | 13.58 |
| Left ILF and Right VOF | 0.31 | 0.13 | 16.18 |
| All seven tracts | 0.345 | 0.84 | 29.32 |
| MTV | | | |
| Left ILF | 0.30 | 0.041 | 13.55 |
| Right ILF | 0.073 | 0.35 | 13.84 |
| Left OR | 0.22 | 0.089 | 13.66 |
| Right OR | 0.13 | 0.20 | 13.77 |
| Forceps major | 0.11 | 0.26 | 13.80 |
| Left VOF | 0.11 | 0.26 | 13.80 |
| Right VOF | 0.33 | 0.033 | 13.52 |
| Left ILF and Right VOF | 0.35 | 0.093 | 16.12 |
| All seven tracts | 0.83 | 0.054 | 27.99 |

**Table S1. The statistical evaluation for linear regression models of stereoacuity using properties of visual tracts.** The left column indicates the representative input variables in the models. The middle two columns indicate goodness-of-fit statistics of the regression models ($R^2$ and p-value of F-test). The right column indicates the Bayesian Information Criterion (BIC) of the regression models (see Materials and Methods). The dark and light gray rows indicate modes that were significant ($p < 0.05$), with the dark gray rows indicating the best model that showed the lowest BIC. The model using only the right VOF showed the lowest BIC in both FA and MTV. While the model using only the left ILF was significant for MTV, the model for FA as well as the group analysis (Fig. 2D) did not support the relationship between left ILF and stereoacuity.
